# Age and Sex-Specific Changes in Mitochondrial Quality Control in Skeletal and Cardiac Muscle

**DOI:** 10.1101/2025.01.15.633257

**Authors:** Catherine B. Springer-Sapp, Olayinka Ogbara, Maria Canellas Da Silva, Yuan Liu, Steven J. Prior, Sarah Kuzmiak-Glancy

## Abstract

Skeletal and cardiac muscle mitochondria exist in a dynamic reticulum that is maintained by a balance of mitochondrial biogenesis, fusion, fission, and mitophagy. This balance is crucial for adequate ATP production, and alterations in skeletal muscle mitochondria have been implicated in aging-associated declines in mitochondrial function. We sought to determine whether age and biological sex affect mitochondrial content [Complex IV (CIV)], biogenesis (PGC-1ɑ), fusion (MFN2, OPA1), fission (DRP1, FIS1), and mitophagy (Parkin, Pink1) markers in skeletal and cardiac muscle by assessing protein expression in tibialis anterior (TA) and ventricular tissue from 16 young (≤6 months) and 16 old (≥20 months) male and female Sprague-Dawley rats. In the TA, CIV expression was 40% lower in old vs. young rats (p<0.001), indicating lower mitochondrial content, and coincided with higher expression of Parkin (+4-fold, p<0.001). Further, MFN2 expression was higher (+2-fold, p<0.005) and Parkin was lower (-40%, p=0.014) in older rats. In cardiac muscle, mitochondrial content was maintained in old vs. young rats, and this occurred concomitantly with higher expression of both PGC-1ɑ and Parkin. MFN2 and OPA1 expression were also 1.2-5-fold higher in older rats (p<0.05 for all). Largely, protein expression did not differ between male and female rats, with the exception of Pink1 and FIS1 expression in the TA. Collectively, older skeletal and cardiac muscle demonstrated higher expression of fusion and mitophagy proteins, which indicates age alters the balance of biogenesis, fission, fusion, and mitophagy. This may, in turn, affect the ability to provide ATP to these metabolically active tissues.

## Introduction

Mitochondrial dysfunction is frequently observed in aging muscle, and previous studies have reported decreases in mitochondrial respiration^1^, reduced time to open the mitochondrial permeability transition pore^1^, and increased mitochondrial DNA damage^2^ in older skeletal and cardiac muscle. These changes coincide with alterations in the structure of mitochondria in old vs. young skeletal and cardiac muscle that may negatively impact the function of muscle mitochondria in aging.

In healthy skeletal and cardiac muscle, mitochondria are organized in a reticulum to efficiently produce and distribute ATP. This reticulum is maintained by multiple processes that are collectively referred to as mitochondrial quality control. Together, the processes of fusion, fission, mitophagy, and biogenesis ensure the mitochondrial reticulum is able to meet the energetic demands of the cell, while mitigating dysfunction. Constant cycles of fusion and fission ensure the mitochondrial reticulum functions optimally by redistributing matrix components and preserving membrane potential^3^. Fusion creates a more expansive reticulum by joining adjacent mitochondria through outer (OMM) and inner mitochondrial membrane (IMM) proteins. OMM-bound proteins mitofusin 1 and 2 (MFN1 and MFN2) allow for OMM fusion, while IMM protein optic atrophy 1 (OPA1) fuses the IMM. Opposing fusion, is the process of fission which separates dysfunctional portions from the reticulum^4^. OMM proteins such as fission 1 (FIS1) and mitochondrial fission factor (MFF) recruit dynamin-related protein 1 (DRP1) to the surface of the mitochondria^5^. DRP1 physically cinches the mitochondrion causing it to cleave into two distinct sections of mitochondria^4^.

Cleaved mitochondria can subsequently undergo selected degradation via mitophagy. The primary mitophagy pathway is the phosphatase and tensin homologue-induced putative kinase 1 (Pink1) and Parkin pathway which occurs on the outer mitochondrial membrane (OMM). The accumulation of Pink1 on the OMM recruits and phosphorylates E3 ubiquitin ligase Parkin and causes ubiquitin chains to form^6^. The ubiquitin chain serves as a binding site for a phagophore, which will engulf the dysfunctional mitochondria before merging with a lysosome and degrading the encapsulated mitochondria. Conversely, mitochondrial biogenesis is regulated by the transcriptional coactivator peroxisome proliferator-activated receptor-gamma coactivator 1 alpha (PGC-1α). PGC-1α translocates to the nucleus and activates several downstream transcription factors to promote the synthesis of new mitochondria^7^. Carefully balanced mitophagy and biogenesis, along with fusion and fission are therefore crucial for quality control and maintaining the mitochondrial reticulum.

Previous investigations have produced interesting, but at times disparate, data regarding how age affects mitochondrial quality control. Some studies provide evidence of reduced fusion, with lower MFN1 and MFN2 protein content in skeletal muscle from old vs. young animals^8–13^, but OPA1 expression appears largely unchanged^8,11,14^. Fission and mitophagy may be increased in older age as FIS1 protein expression was shown to be higher in skeletal muscle of old animals^9–12^, DRP1 protein expression is the same^8,11,15^ or higher^12^ in old vs. young, and the mitophagy-related protein Parkin protein expression increases with age^9,16^. Relatively few studies have addressed cardiac muscle, but MFN1 and MFN2 protein expression have been shown to be lower in old vs. young cardiac muscle^15,17^, while OPA1 protein expression may be increased^18^ and DRP1 protein expression does not appear to change^17^, highlighting some potential differences between skeletal and cardiac muscle.

While some patterns are emerging, the use of different muscles, animal models, and aging timepoints does not allow for a cohesive understanding of how age affects the expression of proteins controlling mitochondrial content (biogenesis and mitophagy) and morphology (fission and fusion). Further, limited data exist in aged cardiac tissue, likely stemming from the previously held notion that fusion and fission are unnecessary for normal heart function^19^, and differences that may exist due to biological sex remains understudied. Notably, only three of eleven aforementioned studies including females, and only one study included a comparison between non-transgenic male and female mice^16^. Triolo et al.^16^ found higher Parkin protein expression in skeletal muscle from old vs. young mice, but also found Parkin expression to be higher in the female compared with male mice. The expression of autophagy-related protein Beclin-1 was also higher in the old animals; however, when sexes were separated, only the old male mice had significantly higher Beclin-1 protein expression than the young group^16^, demonstrating the potential for sex-specific changes with age. Therefore, the purpose of this study was to determine the effects of age and sex on mitochondrial content and the expression of mitochondrial quality control proteins regulating biogenesis, mitophagy, fusion and fission in both cardiac and skeletal muscle (tibialis anterior) of male and female rats.

## Methods

### Animals

All animal research procedures were approved and conducted in accordance with the University of Maryland Institutional Animal Care and Use Committee (R-FEB-23-03). Sixteen young (2-5 months) and 16 old (21-34 months) adult Hilltop SD rats (Rattus norvegicus, Scottdale, PA, USA) were pair-housed fed ad-libitum with standard chow. Groups were further stratified by biological sex resulting in four groups with eight rats in each group. Rats were sacrificed at < 6 months (young) or >20 months of age.

On the day of sacrifice, rats were massed and anaesthetized with 2-4% isoflurane supplemented with 100% oxygen. Rats were placed supine on the surgical area and upon cessation of toe and tail-pinch reflexes, a thoracotomy was performed, and death was induced by exsanguination. Hearts were quickly removed and cannulated via the aorta and rinsed with phosphate buffer saline. Subsequently, the skin on the hindlimbs of the mice was removed, and the tibialis anterior (TA) from both legs were quickly excised. Heart and TA masses were determined before storing at -20°C. Hindlimb lengths (tibial, and knee-to-heel) were measured with digital calipers.

### Protein Expression

Mitochondrial quality control protein content was quantified using western blots. MFN2 and OPA1 were used as markers of mitochondrial fusion, while DRP1 and FIS1 were used as markers of mitochondrial fission. Parkin and Pink1 were used as markers of mitochondrial mitophagy, and mitochondrial biogenesis was measured by PGC-1α protein expression. Finally, mitochondrial content was quantified by CIV protein expression.

Proteins were extracted from tissues in 1% Triton buffer with HALT protease and phosphatase inhibitor cocktails (Thermo Fisher Scientific 78430 and 78428, Waltman, MA, USA). Whole muscle homogenates were made for both the TA and heart using a hand-held homogenizer and subsequently centrifuged at 500 x g for 5 minutes at 4°C. Supernatants were collected and protein concentration was quantified by Pierce bicinchoninic acid assay (Thermo Fisher Scientific 23225, Waltham, MA, USA). Equal amounts of protein (30ug for TA; 60ug for heart) were separated by SDS-PAGE on 4-15% Bio-Rad TGX stain free gels (4568085, Hercules, CA, USA) before photoactivation of gel. Proteins were transferred onto polyvinylidene difluoride (PVDF) membranes (Bio-Rad 1704156, Hercules, CA, USA). Protein transfer was confirmed by imaging total protein fluorescence. Membranes were blocked for 2 hours at room temperature in TBS (Bio-Rad 1706435, Hercules, CA, USA) with 5% milk or BSA (Fisher Scientific BP1600-100, Hampton, NH, USA) and 0.1% Tween-20 (Bio-Rad 1610781, Hercules, CA, USA). Membranes were incubated overnight at 4° C with primary antibodies (Supplemental Table 1). A secondary antibody conjugated to horseradish peroxidase (Supplemental Table 1) was used to detect bands in conjunction with Clarity Western Enhanced Chemiluminescent substrate from Bio-Rad (Hercules, CA, USA). Band intensities were captured using a Bio-Rad ChemiDoc XRS and analyzed using Image Lab 8.0 (Bio-Rad, Hercules, CA, USA). Molecular weights of proteins were confirmed by comparing to a protein standard (Bio-Rad 1610373, Hercules, CA, USA). Band intensities were normalized to total protein in the corresponding lane. In secondary analyses to account for differences in mitochondrial content, MFN2, OPA1, FIS1, DRP1, Parkin, and Pink1 content were expressed relative to complex IV (CIV) content. Representative western blot images are shown in Supplemental Figures 1-4.

### Statistical Analysis

Statistical analyses were performed using GraphPad Prism 9 (GraphPad Software, Boston, MA). Mitochondrial quality control protein expression was analyzed separately for TA and cardiac muscle using two-way ANOVAs with age and sex as factors. Following significant interaction effects, post-hoc comparisons were made using Fisher’s LSD. All tests were two-tailed with statistical significance set at P < 0.05. Data are presented as means ± standard error of the mean.

## Results

### Rat Characteristics

Rat characteristics are summarized in Table 1. Old rats were larger (P <0.001) and had 83% larger hearts and 39% larger TA masses (P < 0.001) compared with young rats; however, old animals had lower skeletal muscle mass relative to body mass than young rats as evident by 31% lower TA to body mass ratio (P < 0.001). Body mass relative to tibial and heel to knee lengths were higher in old animals compared with young counterparts (P < 0.001), likely driven by higher fat mass in the older animals.

**Table 1.** Rat characteristics.

|  | **Young Female**<br>**(n = 8)** | **Young Male**<br>**(n = 8)** | **Old Female**<br>**(n = 8)** | **Old Male**<br>**(n = 8)** |
| --- | --- | --- | --- | --- |
| Age (weeks) | $8.5 \pm 0.2^{* \#}$ | $13.3 \pm 2.0^*$ | $109.9 \pm 10.1$ | $93.5 \pm 2.7$ |
| Body mass (g) | $200 \pm 4.9^{* \#}$ | $331 \pm 24.5^*$ | $393 \pm 18.4^{\#}$ | $697 \pm 46.1$ |
| Heart mass (g) | $0.746 \pm 0.016^{* \#}$ | $1.2 \pm 0.046^*$ | $1.4 \pm 0.078^{\#}$ | $2.1 \pm 0.123$ |
| TA mass (g) | $0.395 \pm 0.006^{* \#}$ | $0.655 \pm 0.047^*$ | $0.547 \pm 0.041^{\#}$ | $0.931 \pm 0.040$ |
| Tibial length (mm) | $34.6 \pm 0.2^{* \#}$ | $37.4 \pm 0.6^*$ | $43.0 \pm 0.4^{\#}$ | $48.4 \pm 1.0$ |
| Heel to knee length (mm) | $43.4 \pm 0.7^{* \#}$ | $47.8 \pm 0.5^*$ | $51.1 \pm 0.3^{\#}$ | $58.5 \pm 0.7$ |
| TA mass/body mass ratio | $0.00195 \pm 0.00004^*$ | $0.00198 \pm 0.00004^*$ | $0.00136 \pm 0.00008$ | $0.00136 \pm 0.00006$ |
| Heart mass/body mass ratio | $0.00369 \pm 0.00008$ | $0.00355 \pm 0.00019$ | $0.00358 \pm 0.00036$ | $0.00311 \pm 0.00019$ |
| Body mass/tibial length ratio | $5.9 \pm 0.1^{* \#}$ | $8.1 \pm 0.1^*$ | $9.2 \pm 0.4^{\#}$ | $14.2 \pm 1.1$ |
| Body mass/heel to knee length | $4.7 \pm 0.1^{* \#}$ | $6.3 \pm 0.2^*$ | $7.7 \pm 0.3^{\#}$ | $11.7 \pm 0.9$ |

Legend = TA, tibialis anterior. Data are means ± SEM. $^*P < 0.05$ different from old, within the same sex; $^{\#}P < 0.05$ different from males, within the same age group.

Overall, young and old female rats were smaller than their male counterparts. Compared with young males, young females had 66% lower body masses (P < 0.001), 54% lower heart masses (P < 0.001), and 66% lower TA masses (P < 0.001). Similarly, old females had 71% lower body masses (P < 0.001), 50% lower heart masses (P < 0.001), 70% lower TA masses (P < 0.001) than old males. Females also had lower body mass to tibial length ratio and body mass to heel to knee lengths when compared with male counterpart of similar age (P < 0.001). Neither young nor old females TA to body mass ratio differed from males, signifying relative skeletal muscle mass was similar between sexes (P = 0.60 and P = 0.99, respectively).

### Skeletal muscle mitochondrial content, mitophagy, biogenesis, and quality control

Mitochondrial content: Skeletal muscle mitochondrial content was lower in old compared to young animals and was likely driven by an increase in mitophagy. Mitochondrial content, as measured by CIV protein expression, was 40% lower in the older TAs compared with the young counterparts (Figure 1A, main effect of age P < 0.001), with no effect of biological sex. While GAPDH was initially going to be used to normalize protein expression, GAPDH protein expression was 1.3-fold higher in the old TAs compared with the young (Figure 1B, main effect of age P < 0.005), making GAPDH inappropriate to use as a loading control. Thus, expression of specific proteins was normalized to total protein in each sample as acquired from stain-free imaging. Additionally, mitochondrial quality control protein expression was also expressed relative to CIV expression to ensure changes in protein expression were not driven by changes in mitochondrial content.

**Figure 1.**
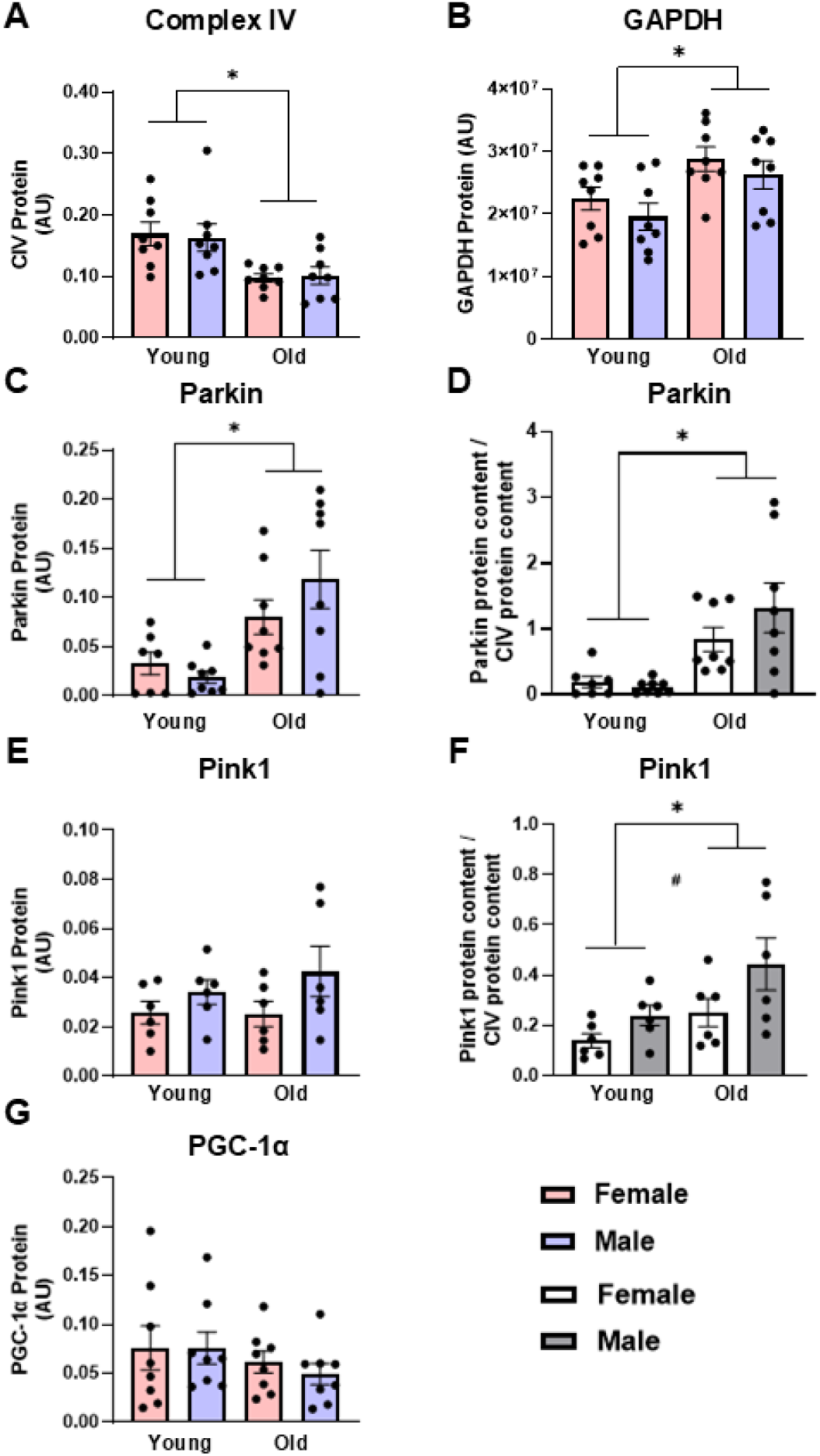
Complex IV (CIV, panel A) protein expression is lower, glyceraldehyde-3-phosphate dehydrogenase (GAPDH, panel B) and Parkin (panels C & D) protein expression are higher, and Pink1 (panels E & F) and peroxisome proliferator-activated receptor gamma coactivator 1-alpha (PGC-1α, panel G) protein expression are not different in old vs. young tibialis anterior muscle. Data in panels A, B, C, E, and G are normalized to total protein. Data in panels D and F are normalized to total protein and subsequently expressed relative to CIV expression. Legend = AU, arbitrary units; YF, young female; YM, young male; OF, old female; OM, old male. #Main effect of sex (P < 0.05), *Main effect of age (P < 0.05). Data are presented as means + SEM, n = 6-8 per group.

Mitophagy and biogenesis: Expression of markers of mitophagy was increased in old skeletal muscle. Parkin protein expression was almost 4-fold higher in old TAs compared with young counterparts (Figure 1C, main effect of age P < 0.001). Pink1 expression did not differ between young and old rat TAs (Figure 1E). When Parkin was expressed relative to CIV, the age differences were further exacerbated, with old rat TAs having 7-fold higher Parkin expression compared with young animals (Figure 1D, main effect of age P <0.001). Pink1 expression was 83% higher in the old animals compared with young (Figure 1F, main effect of age P = 0.023), as well as 75% higher in males compared with to females when Pink1 was expressed relative to CIV (Figure 1F, main effect of sex P = 0.033). PGC-1α expression was measured as a marker of mitochondrial biogenesis but this did not differ between young and old rats (Figure 1G, main effect of age P = 0.211), with no sex-specific changes in PGC-1α protein expression.

Fusion and fission: Mitochondrial fusion, but not fission, was higher in skeletal muscle from old compared with young animals. OMM fusion protein MFN2 and the long and short isoforms of IMM fusion protein OPA1 were analyzed. MFN2 protein expression was 2-fold higher in the TAs from the old animals compared with young animals (Figure 2A, main effect of age P < 0.005). Conversely, OPA1-Short and OPA1-Long protein content were unaffected by age or sex (Figure 2C and 2E, respectively). These results were consistent when MFN2, OPA1-Short, and OPA1-Long protein content were subsequently expressed relative to CIV content, with MFN2 protein levels 3.5-fold higher in old TAs compared with young when normalized to CIV expression (Figure 2B,D,F, main effect of age P < 0.001). No fusion proteins were significantly impacted by sex regardless of CIV normalization.

**Figure 2.**
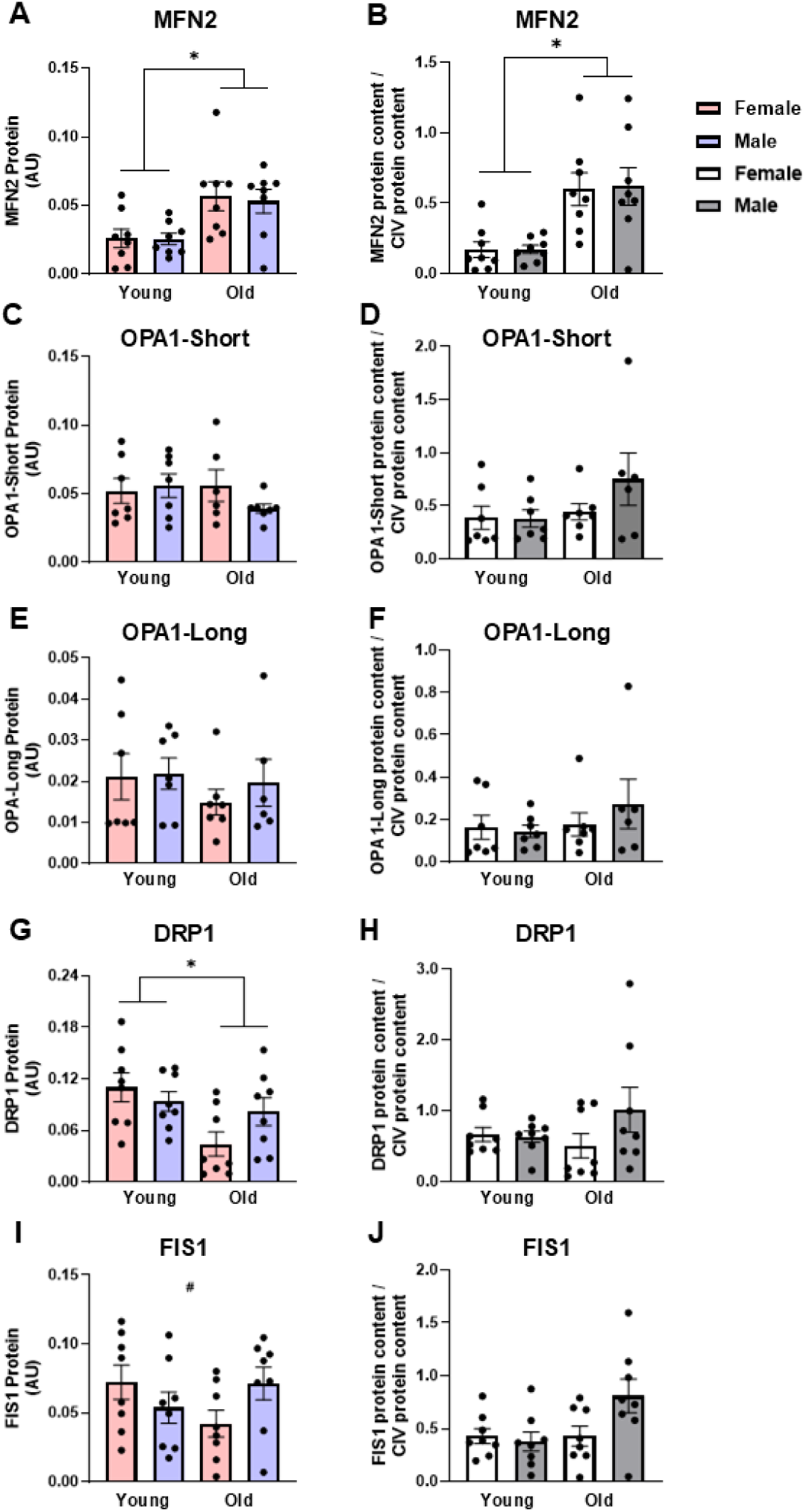
Mitofusin 2 (MFN2, panels A & B) protein expression is higher, optic atrophy 1-short (OPA1-Short, panels C & D) and OPA1-Long (panels E &F) protein expression are not different, dynamin-related protein 1 (DRP1, panels G & H) protein expression is lower in old vs. young tibialis anterior muscle, while there was an age*sex interaction for fission 1 (FIS1, panels I & J) protein expression. Data in panels A, C, E, G, and I are normalized to total protein. Data in panels B, D, F, H, and J are normalized to total protein and subsequently expressed relative to CIV expression. Legend = AU, arbitrary units; YF, young female; YM, young male; OF, old female; OM, old male. *Main effect of age (P < 0.05), #Interaction effect (P < 0.05). Data are presented as means + SEM, n = 7-8 per group.

DRP1 expression was 40% lower in the old rat TAs compared with young (Figure 2G, main effect of age P = 0.014), with no differences between sexes. There was a significant sex * age interaction effect for FIS1 protein expression (Figure 2I, interaction P = 0.047): While FIS1 expression was numerically lower in old vs. young females and numerically higher in old vs. young males, individual group differences did not reach statistical significance. The age and sex-related differences in DRP1 and FIS1 protein expression were likely due to differences in mitochondrial content as these differences were abolished when DRP1 and FIS1 were expressed relative to CIV content (Figure 2H and 2J, respectively).

### Cardiac muscle mitochondrial content, mitophagy, biogenesis, and quality control

Mitochondrial content, mitophagy, and biogenesis: Unlike skeletal muscle, mitochondrial content was maintained in old cardiac muscle, but like skeletal muscle, old animals had higher mitophagy protein expression in the heart. CIV protein expression did not differ between young and old rat cardiac muscle (Figure 3A). Parkin protein expression was over 5-fold higher in the cardiac muscle from old rats compared with young counterparts (Figure 3C, main effect of age P = 0.019) and remained over 5-fold higher in old cardiac muscle when Parkin was expressed relative to CIV expression (Figure 3D, main effect of age P = 0.020). Contrary to findings in skeletal muscle, PGC-1α protein expression in old cardiac muscles was almost 1.5-fold higher than young cardiac muscles (Figiure 3E, main effect of age P < 0.005), indicating a concomitant increase in biogenesis and mitophagy in old cardiac tissue. No sex-specific differences in markers of mitophagy or biogenesis were detected between young and old rat cardiac muscles.

**Figure 3.**
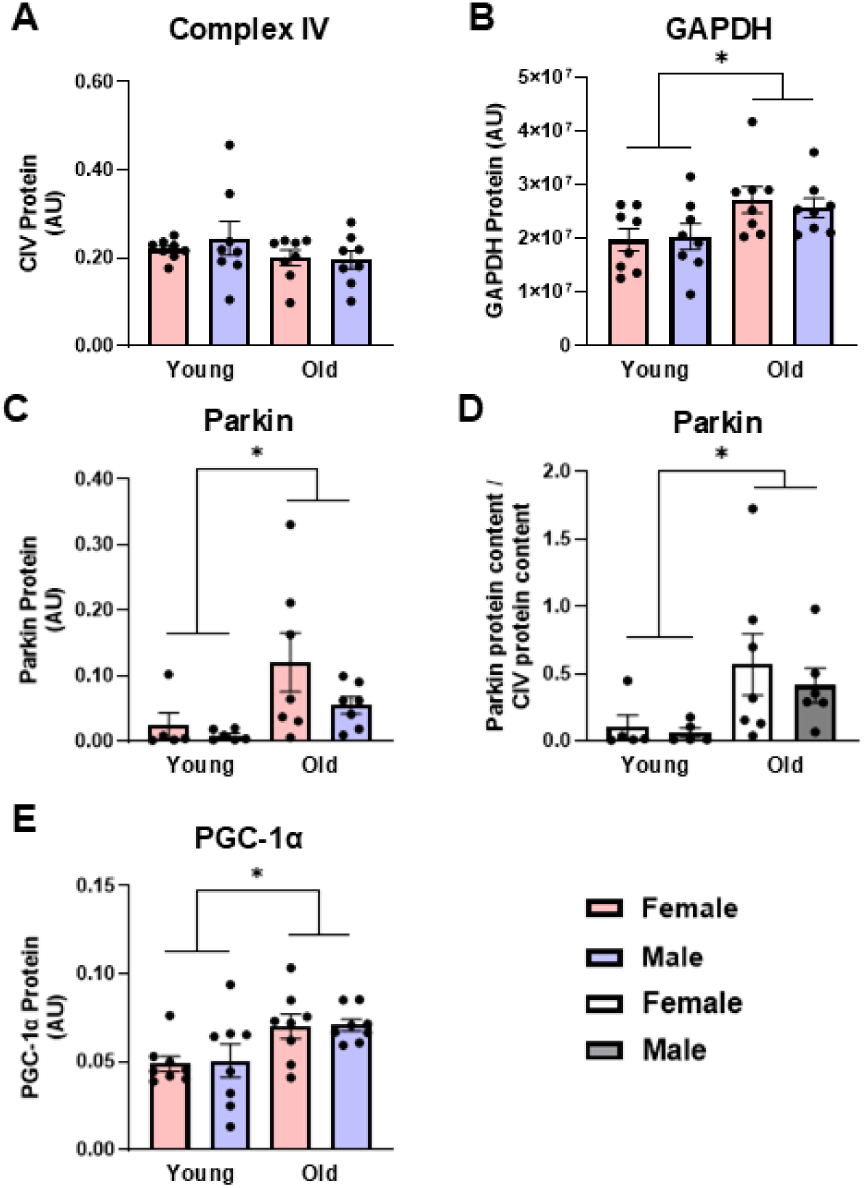
Complex IV (CIV, panel A) protein expression is not different, while glyceraldehyde-3-phosphate dehydrogenase (GAPDH, panel B), Parkin (panels C & D), and peroxisome proliferator-activated receptor gamma coactivator 1-alpha (PGC-1α, panel E) protein expression are higher in old vs. young cardiac muscle. Data in panels A, B, C, and E are normalized to total protein. Data in panel D are normalized to total protein and subsequently expressed relative to CIV expression. Legend = AU, arbitrary units; YF, young female; YM, young male; OF, old female; OM, old male. *Main effect of age (P < 0.05). Data are presented as means ± SEM, n = 5-8 per group.

Fusion and fission: Age significantly impacted fusion protein expression in cardiac muscles, while sex did not. MFN2 was over 2-fold higher in old cardiac muscles compared with young (Figure 4A, main effect of age P = 0.018). Similarly, OPA1-Short and OPA1-Long protein expression were 1.2- and 1.8-fold higher in the cardiac muscle from old rats compared with young, respectively (Figures 4C and 4E, main effects of age P = 0.049 and P < 0.001, respectively). This pattern persisted when MFN2, OPA1-Short and OPA1-Long were expressed relative to CIV expression (Figures 4B, 4D, and 4F, main effect of age P < 0.03 for all). There were no significant differences in cardiac muscle DRP1 or FIS1 protein expression between young or old, male or female animals, indicating no significant effect of age (Figures 4G and 4I, respectively). Expressing DRP1 and FIS1 protein expression relative to CIV expression did not alter these results (Figures 4H and 4J, respectively).

**Figure 4.**
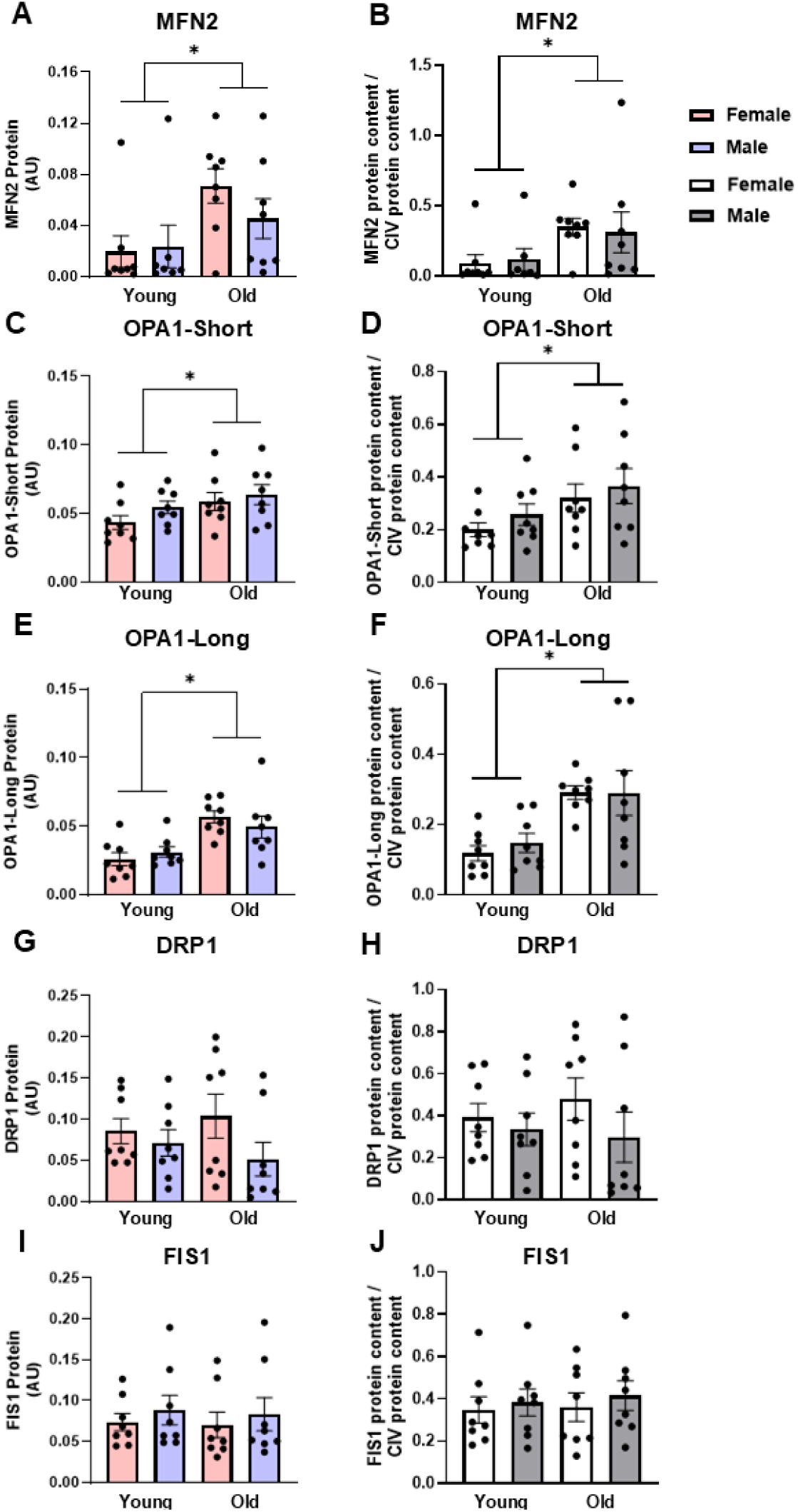
Mitofusin 2 (MFN2, panels A & B), optic atrophy 1-short (OPA1-Short, panels C & D) and OPA1-Long (panels E &F) protein expression are higher, while dynamin-related protein 1 (DRP1, panels G & H) and fission 1 (FIS1, panels I & J) protein expression are not different in old vs. young cardiac muscle. Data in panels A, C, E, G, and I are normalized to total protein. Data in panels B, D, F, H, and J are normalized to total protein and subsequently expressed relative to CIV expression. Legend = AU, arbitrary units; YF, young female; YM, young male; OF, old female; OM, old male. *Main effect of age (P < 0.05). Data are presented as means ± SEM, n = 7-8 per group.

## DISCUSSION

The data in this comprehensive analysis demonstrate that age affects mitochondria from both skeletal and cardiac muscle. While patterns exist in how age affects quality control in both tissues, key differences may underlie how aging differentially affects these tissues. First, old skeletal and cardiac muscle both show markedly higher expression of proteins regulating mitophagy, indicating a need for a greater removal of damaged mitochondria in these tissues in older animals. Interestingly, in skeletal muscle, this increase in mitophagy is not accompanied by an increase in biogenesis proteins and is associated with lower mitochondrial content in old compared with young animals. In cardiac muscle from old animals, there is higher expression of biogenesis proteins in concert with higher mitophagy proteins, and mitochondrial content is maintained. Second, both skeletal and cardiac muscle from old animals have higher expression of proteins regulating outer membrane fusion, while fission proteins are unchanged in cardiac muscle yet decreased in skeletal muscle in old animals. These indicate that aging has differential effects on mitochondrial quality control in skeletal vs. cardiac muscle. Finally, the present investigation shows GAPDH is an unreliable housekeeping/normalization protein for skeletal and cardiac muscle when age is a variable in the investigation. GAPDH expression in TA and cardiac muscle from old rats was 30-40% higher than in young rats. Together, our results demonstrate alterations to expression of mitochondrial quality control proteins with age that differ by muscle type but are largely not different between males and females.

### Skeletal Muscle Mitochondrial Quality Control

The skeletal muscle of old male and female rats have lower mitochondrial content, as measured by CIV protein expression in the TA muscles, compared with their young counterparts.

This is not surprising: studies in mice have report lower CIV protein expression, as well as citrate synthase activity, in old versus young counterparts^10,11^, and microscopy studies show mitochondrial volume density (mitochondria per um^2^) is lower in old compared to young skeletal muscle^15^. Despite the lower mitochondrial content, the present study demonstrates PGC-1α protein expression is not different between young and old groups. This is somewhat surprising, as some studies indicate that consistent with lower mitochondrial content, PGC-1α protein expression is lower in old compared to young skeletal muscle^10,20^, but differences among these and the present study could potentially explain different findings. Interestingly, in one study^20^ PGC-1α was lower in the soleus, but not the plantaris, of older rats, indicating these changes may be muscle- or fiber type-specific. Both investigations studied male rats that were ∼4-14 months older than the rats in the present study, suggesting that a decrease in PGC-1α expression may occur with more advanced age, but taken together with our data, this likely occurs after a decrease in mitochondrial content has already happened.

Consistent with both previous investigations and lower mitochondrial content in old skeletal muscle, mitophagy-related protein expression is higher in old rat skeletal muscle. Parkin protein expression, as well as Pink1 expression normalized to mitochondrial content (CIV), is higher in old rat TA. The accumulation of Pink1 on the OMM leads to the recruitment of Parkin and ultimately mitochondrial degradation, and higher Pink1 protein expression relative to mitochondrial content may indicate more Pink1 accumulation on the OMM due to mitochondrial dysfunction in the old compared with young TA. Indeed, Parkin expression is higher in old compared with young skeletal muscle, consistent with increased mitochondrial degradation in the old animals and previous studies in which Parkin expression is higher in old compared to young mouse quadriceps muscle^16^, and Parkin-independent mitophagy protein Bcl-2 19kD interacting protein (BNIP3) expression is higher in skeletal muscle of old compared with young mice^16^. Further, expression of Parkin and Parkin-independent mitophagy proteins, including BNIP3 and sequestosome 1 (p62), are higher in old male rat extensor digitorum longus and TA muscles^9,21^. Taken together, these data indicate lower mitochondrial content in skeletal muscles of old animals is likely driven by an increase in mitophagy that is not matched with an increase in mitochondrial biogenesis.

Male and female rats did not differ in the majority of aforementioned proteins; however, Parkin expression was higher in male compared with female skeletal muscle, which is consistent with a previous finding from Triolo et al.^16^: that autophagy protein Beclin-1 expression is higher in old male rats than in old females. Given mitophagy is mitochondrial-specific autophagy, it is not surprising that both autophagy- and mitophagy-related protein expression may be upregulated with age. It is unclear why these sex-specific differences are observed, but one hypothesis is that more mitophagy and autophagy in old male rat skeletal muscle might be a consequence of higher ROS production. A recent meta-analysis found ROS measures are higher in skeletal muscle from male subjects across a variety of experimental methods, and specifically, that older men have higher ROS production compared with older women^22^. Elevated levels of ROS can activate both autophagy and mitophagy pathways^23^, therefore it is possible that higher markers of mitophagy in old male rats may be due, in part, to higher ROS.

Intriguingly, while expression of fission proteins DRP1 and FIS1 are not different between young male and young female skeletal muscle, DRP1 protein expression is lower in old rat TAs compared with young and there is a significant sex * age interaction effect for FIS1 protein expression: FIS1 is lower in old compared to young female skeletal muscle, but higher in old compared to young male skeletal muscle. These changes, however, appear to be driven by differences in mitochondrial content, as normalization to CIV protein expression abolished the significance of these effects.

Our results differ from some previous studies that report DRP1 protein expression is the same in old compared with young skeletal muscle^8,11,14,15,24,25^. While the majority of these studies^8,11,15,24,25^ used traditional normalization proteins as loading controls, Leduc-Gaudet et al.^14^ normalized protein expression to total protein via ponceau staining and found DRP1 protein expression is unaltered by age in male mice. Thus, we cannot rule out the possibility that the inclusion of female animals in the present study may drive the significant finding: DRP1 expression was ∼60% lower in old compared with young female skeletal muscle, while it was only ∼15% lower in old vs. young male skeletal muscle and may indicate important differences in the effect of age on mitochondrial fission.

The literature is divided on the effects of age on FIS1 expression, with studies reporting lower^8,13,15^, higher^11,12^, or similar ^24,25^ FIS1 protein expression in old compared with young skeletal muscle. It is unclear why there are such varied results but methodological differences (e.g., animal age, muscle group, protein loading control) along with varied species used (mouse, rat, or human), may account for some of the discrepancies. Indeed, while we found the effect of age on FIS1 protein expression is sex-dependent, these data are no longer significant if normalized to CIV. These conflicting results highlight the need for future studies into sex-specific changes to mitochondrial related proteins with age.

In the present study, MFN2 protein expression is higher in old male and female TA compared with young animals, but neither OPA1-Short nor OPA1-Long was impacted by age or sex. This may indicate greater OMM fusion without a concurrent increase in IMM fusion^26^. Indeed, OPA1 knockdown in young male mice produces enlarged mitochondria with disrupted cristae, demonstrating OMM are able to fuse normally without fully-functioning OPA1^27^. Since fusion is normally regarded as a positive adaptation, higher MFN2 protein expression in aged skeletal muscle may be counterintuitive; but it has also been suggested that increased fusion with age may be a compensatory mechanism to combat mitochondrial dysfunction in skeletal muscle. When mitochondrial membrane potential is disrupted, mitochondria are selected for degradation via mitophagy. In some cases, these dysfunctional mitochondria can fuse with neighboring functional mitochondria as an attempt to rescue membrane potential^26^. The rescuing of mitochondria via fusion can also reduce the proportion of dysfunctional mitochondria^26^. Similarly, fusion allows for the mixing of matrix components, such as mitochondrial DNA (mtDNA), which is thought to reduce the likelihood of mutated mtDNA from reaching pathogenic threshold^28,29^. Diluting mutated mtDNA with normal mtDNA is important for maintaining mitochondrial function as mtDNA encodes for several subunits of the electron transport chain and damage to mtDNA can impact the production and in turn function of these proteins^28,29^. Thus, OMM fusion could be a compensatory mechanism to maintain membrane potential and dilute mutant mtDNA in skeletal muscle. Altogether, these findings suggest a pro-fusion shift in older skeletal muscle, which is consistent with previous studies showing higher expression of fusion relative to fission proteins in older animals^8,13,14^. While the driver of an increase in fusion protein expression remains unclear, greater fusion in old skeletal muscle may indicate a larger need for compensatory fusion due to mitochondrial dysfunction associated with aging.

### Cardiac Muscle Mitochondrial Quality Control

Mitochondrial content does not differ between young and old rat hearts in the present study, and is not surprising considering cardiac tissue relies on constant ATP production and losses in mitochondrial content could have severe consequences^30^. Similar to our findings, others report no changes in CIV protein expression or activity in old compared with young cardiac muscle^17,18^. To account for any numerical, but non-statistical, differences in mitochondrial content, we also expressed mitochondrial quality control proteins in the heart relative to CIV protein. This secondary analysis did not impact any of the results, signifying the differences in mitochondrial quality control proteins were not attributable to differences in mitochondrial content.

Mitochondrial content was likely maintained in old cardiac tissue by higher mitochondrial biogenesis to match the higher mitophagy in old animals. PGC-1α protein expression is higher in old male and female rat hearts compared with young rats, indicating an upregulation of mitochondrial biogenesis. It has previously been reported that PGC-1α gene expression is higher in aged cardiac muscle, likely as a compensatory mechanism to combat mitochondrial dysfunction^31^. The production of new mitochondria likely preserves mitochondrial content in the presence of potentially greater mitochondrial dysfunction and subsequent degradation. Mitochondrial dysfunction in aged cardiac tissue stemming from mutated mtDNA, increased ROS production, and surplus calcium uptake can lead to poor cardiac function with age^30,31^. Mitophagy serves a crucial role in minimizing the negative effects of impaired mitochondrial function. Fittingly, the present study found Parkin expression is higher in old rat hearts compared with young. Limited investigations on mitophagy in aged cardiac muscle, demonstrate higher Parkin and p62 protein content, as well as higher autophagy proteins Beclin-2 and LC3-II in aged mice hearts^32–34^. Together, this indicates mitochondria removal pathways are upregulated in aged cardiac muscle, but in the face of upregulate biogenesis, this likely results in unchanged mitochondrial content.

Relatively few studies have assessed the effects of age on mitochondrial quality control in cardiac muscle; however, abnormal mitochondrial quality control is implicated in multiple cardiovascular adverse events including myocardial ischemia and reperfusion injury, atherosclerosis, and cardiac hypertrophy^35^. This is likely due to the regulatory role mitochondrial structure imparts on function. Indeed, knockout models have demonstrated fusion, fission, and mitophagy proteins (MFN1, MFN2, OPA1, DRP1, Parkin) are necessary for appropriate mitochondrial structure as well as function^36–38^. We found higher protein expression of all markers of fusion (MFN2, OPA1-Short, and OPA1-Long) in old compared with young cardiac muscle, though expression of fission proteins did not differ as a function of age. This pro-fusion shift supports previous findings of enlarged mitochondria in old mouse hearts^39^, and electron microscopy images demonstrating structural changes, but a maintenance of mitochondrial volume in cardiac muscle from old animals^40^. Liang et al.^32^ isolated mitochondria from aged mouse hearts and separated them based on size by differential centrifugation. Large mitochondria had significantly lower Parkin and LC3-II protein expression compared with small mitochondrial from aged hearts^32^. When both mitochondria were pooled together, Parkin and p62 were both elevated in the old hearts^32^. Together, this suggests smaller mitochondria may be targeted for degradation leading to the accumulation of enlarged mitochondria, despite an apparent increase in mitophagy. Such pro-fusion shift in aged hearts may be an adaptive response to inadequate autophagy as an attempt to compensate for mitochondrial damage and increased ROS production^41^.

### Conclusions

Collectively, our results indicate fusion and mitophagy protein expression is higher in aged skeletal and cardiac muscle, while markers of fission are reduced in skeletal muscle of old rats. Similarly, we find that sex interacts with the effect of age on select fission and mitophagy proteins in skeletal, but not cardiac, muscle. Greater fusion and mitophagy may serve as a compensatory mechanism to mitigate mitochondrial dysfunction with age by restoring membrane potential via fusion and degrading unsalvageable mitochondria via mitophagy. In old skeletal muscle, evidence of higher mitophagy without a subsequent increase in biogenesis occurs in tandem with lower mitochondrial content; however, in cardiac muscle the higher mitophagy appears to be offset by biogenesis, maintaining mitochondrial content. These findings underscore key similarities and differences in mitochondrial quality control between young and old skeletal and cardiac muscle that may contribute to differential regulation of mitochondrial morphology and function in different types of muscle with aging.

## CONFLICT OF INTEREST

The authors have no conflicts of interest to declare.

## FUNDING

This work was supported by the National Institute of Aging at the National Institutes of Health (grant number AG064571 to SJP) and the American Heart Association (grant number 16SDG30770015 to SKG).

## ACKNOWLEDGEMENTS & AUTHOR CONTRIBUTIONS

Conception or design of the work: Catherine B. Springer-Sapp, Steven J. Prior, Sarah Kuzmiak-Glancy. Acquisition, analysis, or interpretation of data for the work: Catherine B. Springer-Sapp, Olayinka Ogbara, Maria Canellas Da Silva, Yuan Liu, Steven J. Prior, Sarah Kuzmiak-Glancy. Drafting of the manuscript: Catherine B. Springer-Sapp, Steven J. Prior, Sarah Kuzmiak-Glancy. All authors critically reviewed and approved the final version of the manuscript and agree to be accountable for all aspects of the work. All persons designated as authors qualify for authorship and all those who qualify for authorship are listed.

## Notes

### Competing Interest Statement

The authors have declared no competing interest.

