## Supplemental Table 1; Supplemental Figures 1-4 for "Age and Sex-Specific Changes in Mitochondrial Quality Control in Skeletal and Cardiac Muscle"

**Supplemental Table 1. Antibodies**

| Antibody | Manufacturer | CAS number | Research Resource Identifiers | Dilution |
| --- | --- | --- | --- | --- |
| Anti-rabbit IgG HRP linked antibody | Cell Signaling Technology, Danvers, MA, USA | 7074S | AB_2099233 | 1:2000 in TBS-T with 5% BSA or milk |
| CIV (3E11) Rabbit mAb | Cell Signaling Technology, Danvers, MA, USA | 4850S | AB_2085424 | 1:1000 in TBS-T with 5% BSA |
| DRP1 (D6C7) mAb | Cell Signaling Technology, Danvers, MA, USA | 8570S | AB_10950498 | 1:1000 in TBS-T with 5% BSA |
| Fis1 pAb | Thermo Fisher Scientific, Waltham, MA, USA | PA5-22142 | AB_11152577 | 1:1000 in TBS-T with 5% BSA |
| GAPDH pAb | Thermo Fisher Scientific, Waltham, MA, USA | PA1-987 | AB_2107311 | 1:5000 in TBS-T with 5% BSA |
| Mitofusin-2 Rabbit mAb | Cell Signaling Technology, Danvers, MA, USA | 83667S | AB_2800025 | 1:1000 in TBS-T with 5% milk |
| OPA1 (D6U6N) Rabbit mAb | Cell Signaling Technology, Danvers, MA, USA | 80471S | AB_2734117 | 1:1000 in TBS-T with 5% BSA |
| Parkin Recombinant Rabbit mAb (21H24L9) | Thermo Fisher Scientific, Waltham, MA, USA | 702785 | AB_2842252 | 1:500 in TBS-T with 5% BSA |
| PGC-1α pAb | Thermo Fisher Scientific, Waltham, MA, USA | PA5-72948 | AB_2718802 | 1:500 in TBS-T with 5% milk |
| Pink1 Rabbit mAb | Thermo Fisher Scientific, Waltham, MA, USA | PA1-16604 | AB_2164267 | 1:500 in TBS-T with 5% BSA |

CAS: Catalog Number; IgG: Immunoglobulin G; HRP: Horseradish Peroxidase; mAb: monoclonal; pAb: polyclonal; TBS-T: tris buffer saline with 0.1% Tween 20; BSA: bovine serum albumin.

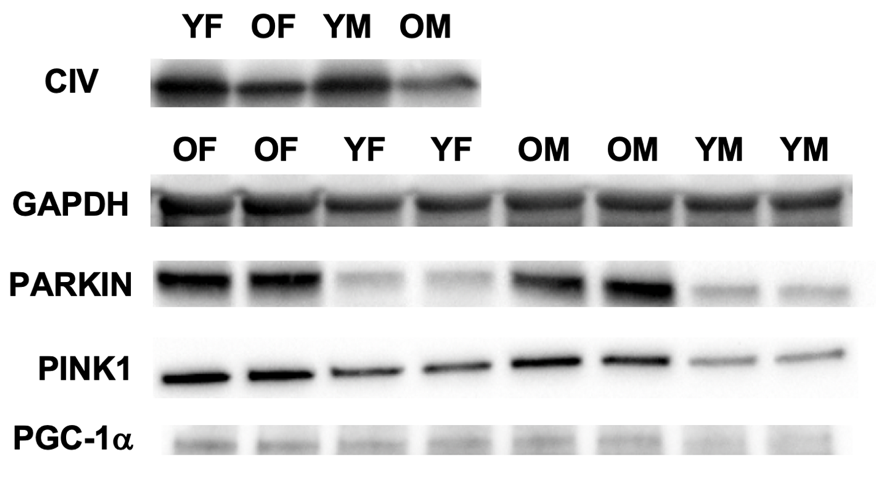
**Supplemental Figure 1**. Representative western blots for markers of mitochondrial content, mitophagy, and biogenesis in tibialis anterior skeletal muscle. YF: Young, female rats; OF: Old, female rats; YM: Young, male rats; OM, Old, male rats.

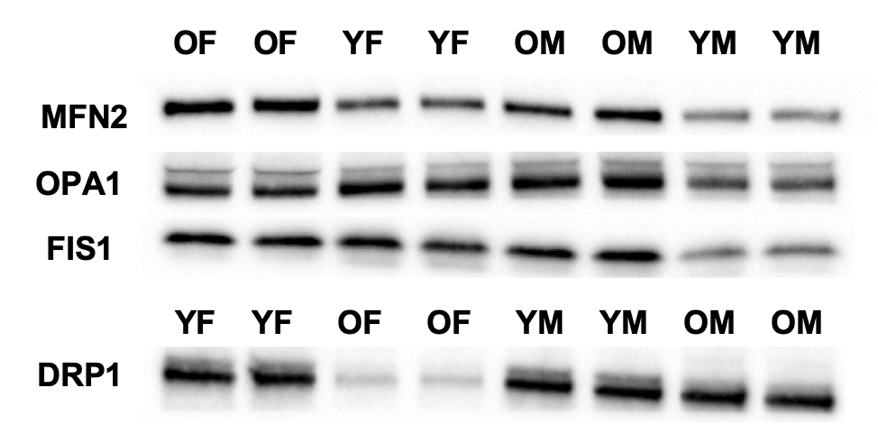

**Supplemental Figure 2**. Representative western blots for markers of mitochondrial fusion and fission in in tibialis anterior skeletal muscle. YF: Young, female rats; OF: Old, female rats; YM: Young, male rats; OM, Old, male rats.

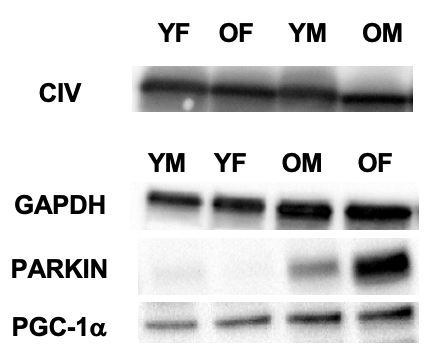

**Supplemental Figure 3**. Representative western blots for markers of mitochondrial content, mitophagy, and biogenesis in cardiac muscle. YF: Young, female rats; OF: Old, female rats; YM: Young, male rats; OM, Old, male rats.

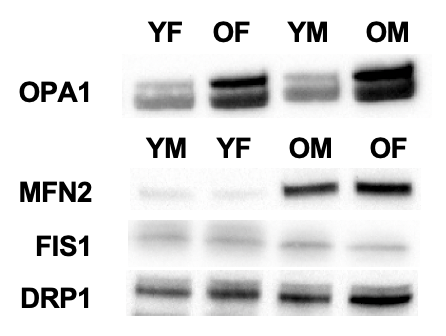

**Supplemental Figure 4**. Representative western blots for markers of mitochondrial fusion and fission in cardiac muscle. YF: Young, female rats; OF: Old, female rats; YM: Young, male rats; OM, Old, male rats.
